# Molecular dynamics simulations reveal the impact of Ser295 phosphorylation on the structure of pyrin domain-containing NOD-like receptor 3

**DOI:** 10.1101/2025.05.28.656610

**Authors:** Christina F. Sandall, Justin A. MacDonald

**Affiliations:** Department of Biochemistry & Molecular Biology, Cumming School of Medicine, University of Calgary, Calgary, Alberta, Canada

**Keywords:** phosphorylation, NACHT domain, nucleotide-binding domain (NBD), inflammasome, electrostatic interactions, molecular dynamics simulations, ATPase

## Abstract

The nucleotide-binding, leucine rich repeat, and pyrin-containing 3 (NLRP3) protein is regulated by phosphorylation of Ser295 in the NACHT domain. This post-translational modification is known to inhibit the enzymatic ATPase activity of NLRP3 and impede inflammasome complex assembly. In this study, modeled structures of unphosphorylated and pSer295-phosphorylated NLRP3-ΔPYD were subjected to molecular dynamics simulations. The outputs showed Ser295 phosphorylation to induce topologic distention of subdomains that comprise the NACHT domain. The relative orientation of important residues within the nucleotide-binding domain (NBD) were altered. Notable structural changes were observed for important residues within the Walker B motif that immediately follow pSer295. A favorable electrostatic environment was created for two residues (Lys232 and His522) that interact with ADP. Several other basic residues could establish favourable charge-charge interactions with the dianionic phosphate of pSer295. Arg296 and Glu343 underwent a functional change from negative/stabilizing to positive/destabilizing interaction upon phosphorylation of Ser295. Taken together, the results suggest that local structural transformations within the NBD could have consequences on the catalytic efficiency of the enzyme and suppress nucleotide turnover.

## INTRODUCTION

The NOD-like receptor containing pyrin 3 (NLRP3) protein is the most comprehensively studied and the prototypical inflammasome-forming member of the family ^1-3^. The NLRP3 protein is comprised of three core functional domains: an N-terminal pyrin domain (PYD), a central nucleotide-binding oligomerization domain (NOD/NBD/NACHT) aligned with the signal transduction ATPases with numerous domains (STAND) class of AAA+ ATPases, and a C-terminal region containing multiple leucine-rich repeat (LRR) domains. As a molecular scaffold, NLRP3 recruits the inflammasome adaptor apoptosis-associated speck-like protein containing a caspase recruitment domain (ASC) protein through homotypic PYD-PYD interactions; this complex is then able to recruit and activate pro-caspase-1 to drive the maturation of pro-inflammatory cytokines (i.e., IL-1β and IL-18) as well as lytic cell death ^1,4^ Transcriptional “priming” is linked to the up-regulation of *NLRP3* signaling ^5^; however, the NLRP3 protein is also subject to a variety of post-translational modifications that are known to “license” the stability, intracellular location, signaling partnerships, and ATPase activity of the protein ^6,7^. Protein phosphorylation offers a particularly effective mechanism to trigger conformational transitions and structural rearrangements of the protein backbone ^8,9^, thereby affecting protein function and regulating signaling pathways. Several phosphorylation sites have been identified for NLRP3 ^10-12^, but the phosphorylation of the Ser295 residue is unique in its potential to modulate the intrinsic ATPase activity of the NACHT domain ^13^.

The ATP-dependency of NLRP3 were first revealed by Duncan and colleagues; mutation of the Walker A motif within the NACHT domain abrogated both *in vitro* ATP-binding and inflammasome-mediated IL-1β secretion from THP-1 cells ^14^. A functional NACHT domain was also required for assembly of the NLRP3 inflammasome complex ^15^. Furthermore, the oligomerization of a phosphomimetic NLRP3 protein (i.e., Ser291Asp, mouse isoform numbering) was abolished in a reconstitution system, whereas the oligomerization of Ser291Ala was unaffected ^16^. The large size and composite domain structure of NLRP3 enables the protein to effectively nucleate oligomeric signaling complexes, thereby recruiting effectors and activating downstream immunological defences. The NACHT domain is thought enable NLRP3 to act as a mechano-chemical coupler –using the energy of ATP hydrolysis to drive global reorganization of protein conformation and provide surface exposure of previously concealed binding sites for oligomeric complex formation ^17,18^.

Since several motifs present in or associated with the NACHT domain play key functional roles in the regulation of substrate binding and catalysis, we postulated that PKA-dependent phosphorylation of Ser295 could result in conformational fluctuations within these regions. In the absence of empirical structures for the phosphorylated form of NLRP3, molecular dynamics (MD) simulations currently provide the only reasonable opportunity to study the effects of Ser295 phosphorylation on the topology of the ATP-binding pocket. This study develops an MD simulation framework to explore the influence of Ser295 phosphorylation on the conformational stability of the NACHT domain.

## METHODS

### MD Simulations

Molecular dynamic (MD) simulations were run with GROMACS v2020.4 ^19^ on a previously optimized model of ADP-bound NLRP3 ^17^. The starting structure employed human NLRP3 (PDB: 6NPY – Chain A) including residues 135-1036, with removal of the N-terminal PYD domain (ΔPYD), bound to ADP ^20^. A dianionic phosphorylated serine residue was substituted for Ser295 in the NLRP3 model using the CHARMM-GUI input generator for PDB file modification ^21^. MD systems were constructed for the unphosphorylated and phosphorylated NLRP3 models with application of the refined all-atom additive CHARMM36m protein force field ^22,23^. System energies were minimized with the steepest descent method, and system equilibration was completed using constant-substance, constant-volume, constant temperature ensemble (NVT) then constant-temperature, constant-pressure ensemble (NPT) for 1000 ps with 1 fs integration time steps. MD production runs were completed as described previously ^17^. GROMOS clustering was performed ^24^, and the medoid structure of the largest cluster observed during simulations was used for the subsequent electrostatic energy calculations.

### Charge-charge Interactions Calculations

The FoldX Suite ^25^ was used to compare the electrostatic energies ⟨Wi⟩ in Ser295-phosphorylated and unphosphorylated NLRP3 due to ion– ion interactions between charged residues with the rest of the ionizable groups in the protein. The total charge–charge interaction energy (⟨Wq-q⟩) for each of the two proteins was calculated as previously described ^26^. Residue interaction networks were defined using the RINGv2.0 webserver ^27^. Covalent and non-covalent bonding events were categorized as: hydrogen bonds if the distance between acceptor and donor atoms was ≤ 3.5 Å and the hydrogen-donor-acceptor angle was ≤ 63°; van der Waals interactions for residues with distances between atom surfaces of ≤ 0.5 Å; or salt-bridges if the distance between the mass centers of oppositely charged groups was ≤ 4.0 Å. Inter-residue distances were calculated with the DISTEVAL webserver ^28^.

### Software

GROMACS, FoldX and ChimeraX were obtained with an academic license. Structural figures were generated with academic versions of the Pymol Molecular Graphics System, the UCSF Chimera Visualization System, Visual Molecular Dynamics (VMD), Avogadro Molecular Editor and the Swiss-PdbViewer (DeepView). Statistical analyses were run in R or GraphPad-Prism. Illustrations were generated using Microsoft PowerPoint, GraphPad-Prism or Adobe Photoshop.

## RESULTS & DISCUSSION

To evaluate the impact of phosphorylation on the topology of the catalytic pocket and the structural motifs that comprise the NACHT domain, a dianionic phosphoryl group was introduced at the Ser295 residue within the ADP-bound NLRP3 model using the CHARMM-GUI input generator for PDB file modification. The modeled structures of phosphorylated and unphosphorylated NLRP3-ADP were subjected to 10 ns molecular dynamics simulations. The two systems were well-behaved with both the unphosphorylated and phosphorylated forms exhibiting stable trajectories before the end of the production runs. Gross comparison of the final equilibrated structures (Figure **1A** and **1B**) suggests Ser295 phosphorylation could induce topologic distention of subdomains that comprise the greater NACHT structure, namely the fish-specific NACHT-associated (FISNA) domain, the winged-helix domain (WHD) as well as helical domain 2 (HD2).

**Figure 1.**
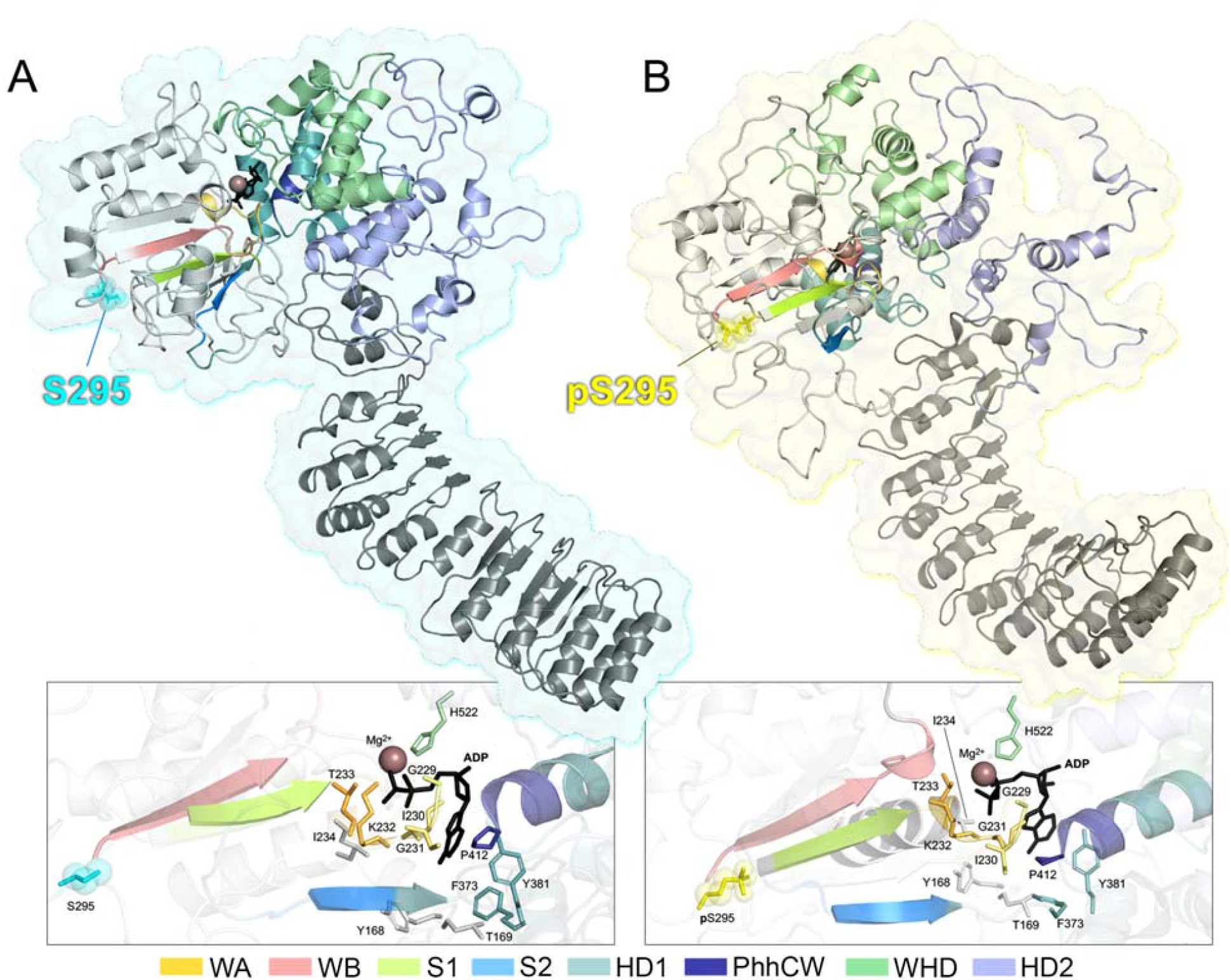
Predominant Medoid Structures Derived from MD Simulations for the Unphosphorylated and pSer295-Phosphorylated Models of NLRP3. Representative structures were selected from the highest occupied cluster over the course of the 10 ns simulation following GROMOS clustering. Cartoon representations of the NLRP3-ΔPYD structure are coloured according to domain and motif; models begin at residue Asp135 and end at the NLRP3 C-terminus (Trp1036). Surface renderings are coloured in cyan (**A**: Ser295, unphosphorylated) or yellow (**B**: pSer295, phosphorylated). Detailed views of ADP-interacting residues and critical secondary structure motifs are provided for the nucleotide-binding pocket of unphosphorylated and phosphorylated NLRP3 to illustrate the conformational and positional changes of key motifs upon phosphorylation of Ser295.

More nuanced changes in the relative orientation of the important residues within the nucleotide-binding domain (NBD) were also apparent with phosphorylation. Using information from NLRP3 structures deposited in the PDB (i.e, 6NPY, 7ALV, 7PZC, and 7VTP), the specific coordination of the ADP nucleotide in the NBD is achieved with hydrogen bonding from Tyr168, Thr169, Gly229, Ile230, Gly231, Lys232, Thr233, Ile234, Tyr381; hydrophobic interactions with Ile234 and Pro412; salt bridges with Lys232 and His522; and π stacking with Phe373. Most of these residues display marked conformational changes in the pSer295 model that would negatively impact nucleotide binding. For example, there is rearrangement of the hydrophobic residues (Ile234, Phe373 and Pro412) responsible for interacting with the adenine base of ADP to stabilise the protein-nucleotide interface. The positioning of Lys232, Thr233, Ile234 as well as His522 are also shifted to impede effective orientation of important side chains that would normally coordinate with the α- and β-phosphates of ADP.

The final phosphorylated and unphosphorylated structure medoids derived from the MD simulations were aligned with ChimeraX, resulting in an RMSD of 5.168 Å and a Q-score of 0.043 over all residues. These outputs suggest a large shift in global protein conformation was elicited by Ser295 phosphorylation (**Figure 2A,B**). Pairwise RMSD analysis suggests the occurrence of large shifts in fold conformation and residue position for several critical functional motifs within the NACHT domain (**Figure 2C**). Notable alterations were observed in the Walker B motif that immediately follows the pSer295 residue. The Walker B motif contains conserved acid residues (Asp302,Asp305) that serve to coordinate and stabilize a Mg2+ ion, which is a mandatory cofactor for the hydrolysis of ATP. The introduction of pSer295 also resulted in higher RMSDs for key residues of the Walker A as well as the Sensor 1 and 2 regions. The Walker A motif represents the “P-loop” in AAA+ ATPases responsible for binding phosphate groups of the nucleotide. Moreover, Sensor 1 contains the critical Arg351 residue that coordinates with the Walker B motif to enable ATP hydrolysis ^15^.

**Figure 2.**
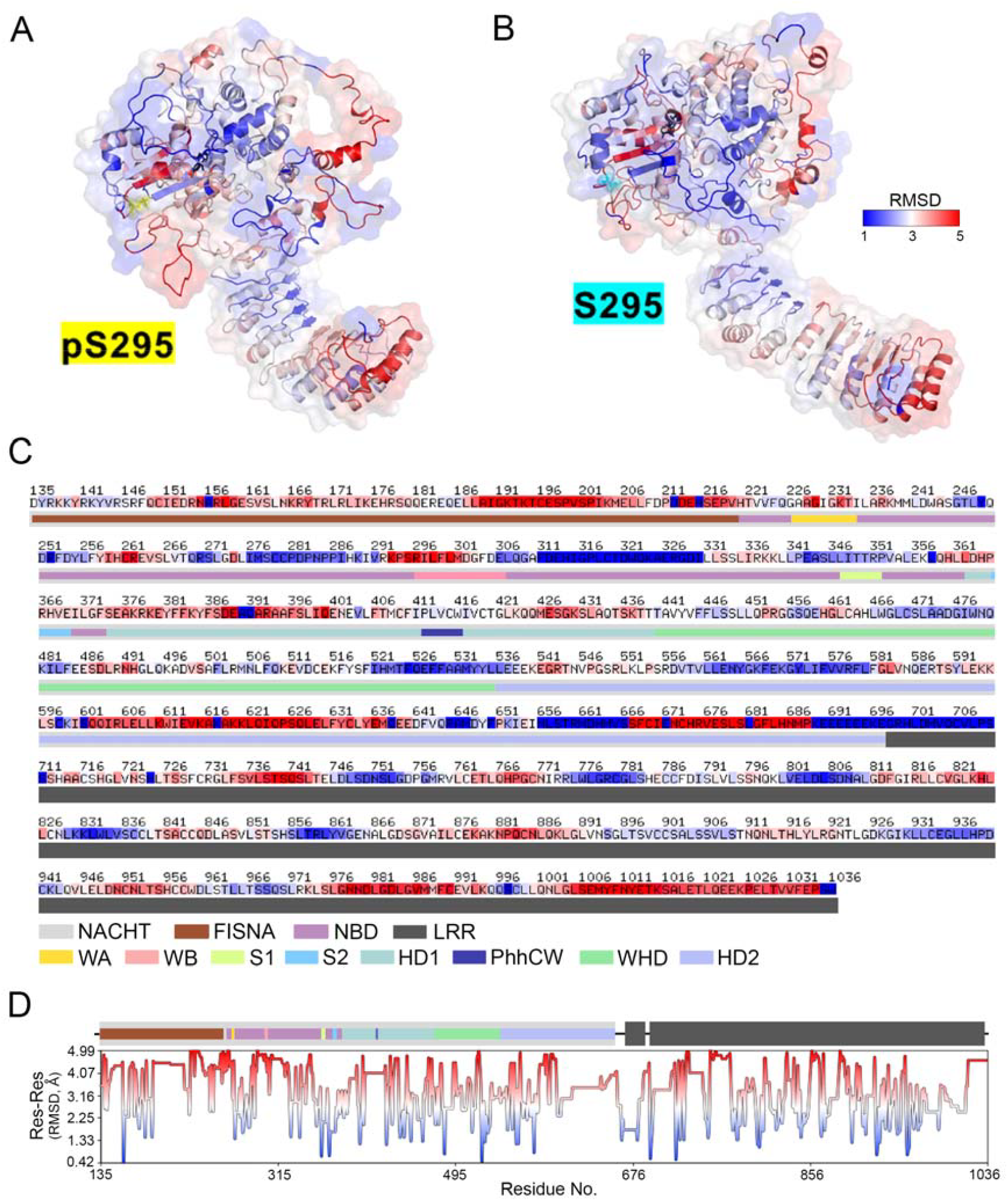
Ser295 phosphorylation elicits a large shift in global NLRP3 protein conformation. Medoids generated from MD simulations of NLRP3 were aligned; representations of the Ser295-unphosphorylated (**A**) and pSer295-phosphorylated (**B**) structures and the primary sequence (**C**) are coloured by individual RMSDs in a spectrum from blue to white to red (representing RMSD values from 0.418 to 4.984 Å). Blue indicates close structural alignment, while higher deviations are in red. In (**D**), residue-residue RMSD values are charted below the sequence. Core domains and motifs of NLRP3 are coloured as indicated: NACHT, domain present in NAIP, CIITA, HET-E, and TP1; FISNA, Fish specific NACHT-associated; NBD, nucleotide-binding domain; LRR, leucine-rich repeat; WA, Walker A; WB, Walker B; S1, Sensor 1; S2, Sensor 2; HD1, Helical domain 1; PhhCW, Phe-(hydrophobic)_2_-Cys-Trp motif; WHD, Winged-helix domain; and HD2, Helical domain 2.

Pronounced structural changes were also observed in the fish-specific NACHT-associated (FISNA) domain that comprises the N-terminal region of NACHT. Xiao and colleagues previously reported that the FISNA domain becomes highly ordered in key regions to stabilize the active NACHT conformation ^29^, namely loop 1 (residues 151-163), helix 2 (residues 176-202), and loop 2 (residues 212-217). Collectively, these regions of the FISNA domain exhibited even more disorder in the pSer295-NLRP3 model, with significant degeneration of the folded conformation (**Figure 1**). Marked alterations in RMSD were noted for the helical domain 1 (HD1) and helical domain 2 (HD2) regions (**Figure 2A,B**). An ordered FISNA domain partners with HD1 and a section of the winged-helical domain (WHD) to stabilize the active NLRP3 oligomerization state ^2,29^. Interestingly, there was minimal impact of Ser295 phosphorylation on the conformation of the winged helix domain (WHD) that is involved in the interaction with ATP, a crucial step in NLRP3 activation. Mutations in the WHD, particularly the conserved His522 residue, can affect ATP binding and hydrolysis ^2^.

Of the 902 residues (i.e., Asp135-Trp1036) present in the models, ionizable side chains (i.e., arginine, lysine, aspartic acid, glutamic acid, histidine, cysteine and tyrosine) accounted for 301 of the 902 amino acids. The energies of individual charge–charge interactions ⟨Wi⟩ for all ionizable residues as compared with all other charged residues were calculated with the FoldX force field that was developed to evaluate the effect of mutations on the stability, folding and dynamic motions of proteins ^25^. The majority of ⟨Wi⟩ values for both phosphorylated (pSer295) and unphosphorylated (Ser295) models are unsurprisingly negative (**Figure 3A**: pSer295, 94.7%; Ser295, 87.7%), indicating favourable interactions with residues of the opposite charge. An approximation of the contribution of charge-charge interactions to the unfolding Gibbs energies (−ΔGq-q) was calculated as ⟨Wq-q⟩ for each model. The ⟨Wq-q⟩ values were negative for both enzyme forms (−306.0 kJ/mol and -328.3 kJ/mol for the unphosphorylated and phosphorylated models, respectively), but indicate that the inclusion of pSer provided more thermodynamic stability. Significant contributors to the Δ⟨Wq-q⟩ of -22.3 kJ/mol between the Ser295 and pSer295 models are labeled in **Figure 3B**, where positive energy values indicate favourable interactions that form in the phosphorylated NLRP3 model, and negative values indicate unfavourable encounters produced upon the phosphorylation of Ser295. These data suggest that a collection of favourable electrostatic compensations induced by the phosphorylation event were mostly localized to residues in close spacial proximity to pSer295 (**Figure 3B**). Additionally, some residues with larger energy differences were located near to the ADP molecule in the catalytic pocket of the NACHT domain; these include residues that promote the nucleotide binding (Walker A) and hydrolysis (Walker B) events that are critical for inflammasome assembly. Nevertheless, several ionizable residues in regions more removed from the either the nucleotide-binding site or the pSer295 residue were also impacted. Taken together, the conformational impact of Ser295 phosphorylation appeared to be transmitted to more remote regions of the protein structure.

**Figure 3.**
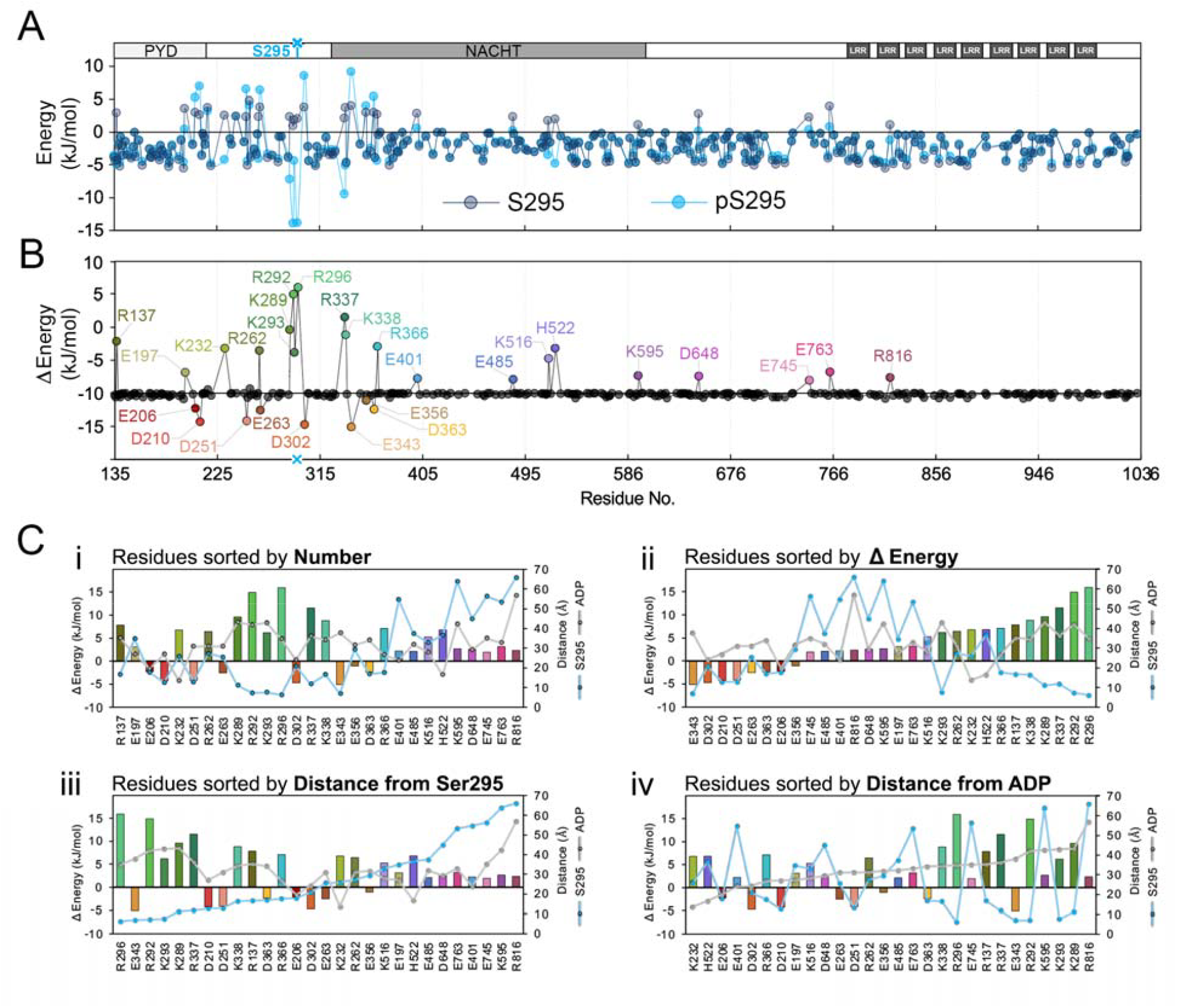
Charge-Charge Interaction Energies for Ionizable Residues in Unphosphorylated (Ser295) and Phosphorylated (pSer295) NLRP3 Models. In (**A**), charge-charge interaction (⟨Wi⟩) energies were calculated with FoldX for all ionizable residues of unphosphorylated (Ser295) and phosphorylated (pSer295) NLRP3. Positive values indicate predominant involvement in destabilizing interactions with like-charged residues, while a negative value indicates contribution to stabilizing interactions with groups of the opposite charge. In (**B**), the residues with the largest contributions to the unfolding Gibbs energies Δ⟨Wq-q⟩ between Ser295- and pSer295-NLRP3 models are plotted. Residues are uniquely coloured for reference. Positive values indicate the development of more favorable interactions for residues in pS295-NLRP3. In (**C**), residues with significant Δ⟨Wi⟩ contributions are collated by: (**i**) primary Residue No.; (**ii**) Δ⟨Wi⟩ energy in ascending order; (**iii**) distance from the Ser295 residue; and (**iv**) distance from the ADP molecule.

The noteworthy residual contributors to the Δ⟨Wq-q⟩ between the Ser295 and pSer295 forms of NLRP3 and their distances from the Ser295 residue and ADP molecule are shown in **Figure 3C,i**.The residues with significant contributions to the overall Δ⟨Wq-q⟩ between models were ranked in ascending order (**Figure 3C,ii**). The top ten residual contributors with favourable contributions to the change in overall conformational energy (i.e., Δ⟨Wq-q⟩) between models included: Arg296, Arg292, Arg337, Lys289 and Lys338. In the case of the five basic residues Arg296, Arg292, Arg337, Lys289 and Lys338, the favourability of interactions can be explained by their distances from Ser295. These five residues reside in close proximity to Ser295, and it is likely that one or several of these positively charged residues establish favourable interactions with the neighbouring dianionic phosphate of pSer295 (**Figure 3D,iii)**. As ADP-bound NLRP3 is known to be inactive and unable to nucleate inflammasome assembly ^2^, it is noteworthy that phosphorylation results in more a favorable electrostatic environment for two residues Lys232 and His522 (**Figure 3D,iv)**. The ε-amino group of Lys232 and the τ-nitrogen of the His522 imidazole group form salt bridges with the β-phosphate group of ADP, with additional bridging to the ⍰-phosphate also provided by His522.

The residues accounting for unfavourable ionic interactions were more dispersed, further reinforcing the hypothesis that the Ser295 phosphorylation event results in structural impacts that can be transmitted globally across the structure. However, Glu343, Asp302, Asp210, Asp251 and Glu263 represent the residues with the most destabilizing charge-charge interactions induced by pSer295 phosphorylation. It is not unexpected that all destabilizing residues are acidic as these would most likely conflict with the introduced dianionic phosphoryl group of pSer295. Moreover, these residues all reside within the NBD region that coordinates nucleotide occupancy. As previously noted, the Asp302 residue resides in the Walker B motif and coordinates binding of the obligate Mg2+ cofactor. Electrostatic destabilization of this critical residue would be expected to provide catalytic inactivation. The relative proximity of Glu343, Asp210 and Asp251 to the ADP molecule (**Figure 3D,iv**) could explain why these destabilizing interactions are observed in the phosphorylated and inactive structure. Alternatively, favourable basic residues involved in coordinating the phosphorylated Ser295 could lose capacity to interact with the unfavourable acidic residues.

To further interrogate these possibilities, the Residue Interaction Network Generator (RINGv2.0) was used to map all the inter-residue bonding distances between all significant contributors to the Δ⟨Wq-q⟩ in the unphosphorylated model structure ^27^. The predicted interactions output (**Figure 4**) suggests Glu343 was within the distance cut-off to hydrogen bond with Arg292, moreover Arg292 was also within the cut-off to bond with Ser295. Thus, in conjunction with the Δ⟨Wq-q⟩ data (**Figure 3**), it is likely that Glu343 underwent an important functional change from negative/stabilizing to positive/destabilizing interaction upon phosphorylation of Ser295 due to a loss of hydrogen-bonding capacity for Arg292. A similar scenario likely occurs for the Arg296 and Asp251 pairing; Asp251 is highly destabilizing, while Arg296 is highly stabilizing upon phosphorylation of Ser295. The change suggests a loss of bonding capacity of Arg296 for Asp251and a shift to Arg296 interaction with pSer295. As a caveat, the residue-residue distances were calculated as an average of all atoms in individual residues, and thus individual atoms may be closer and within interaction cut-offs. The RING outputs predict interactions based on distance cut-offs and atom identities; however, the algorithm does not account for additional proximal residues that provide or interfere with other interactions ^27^.

**Figure 4.**
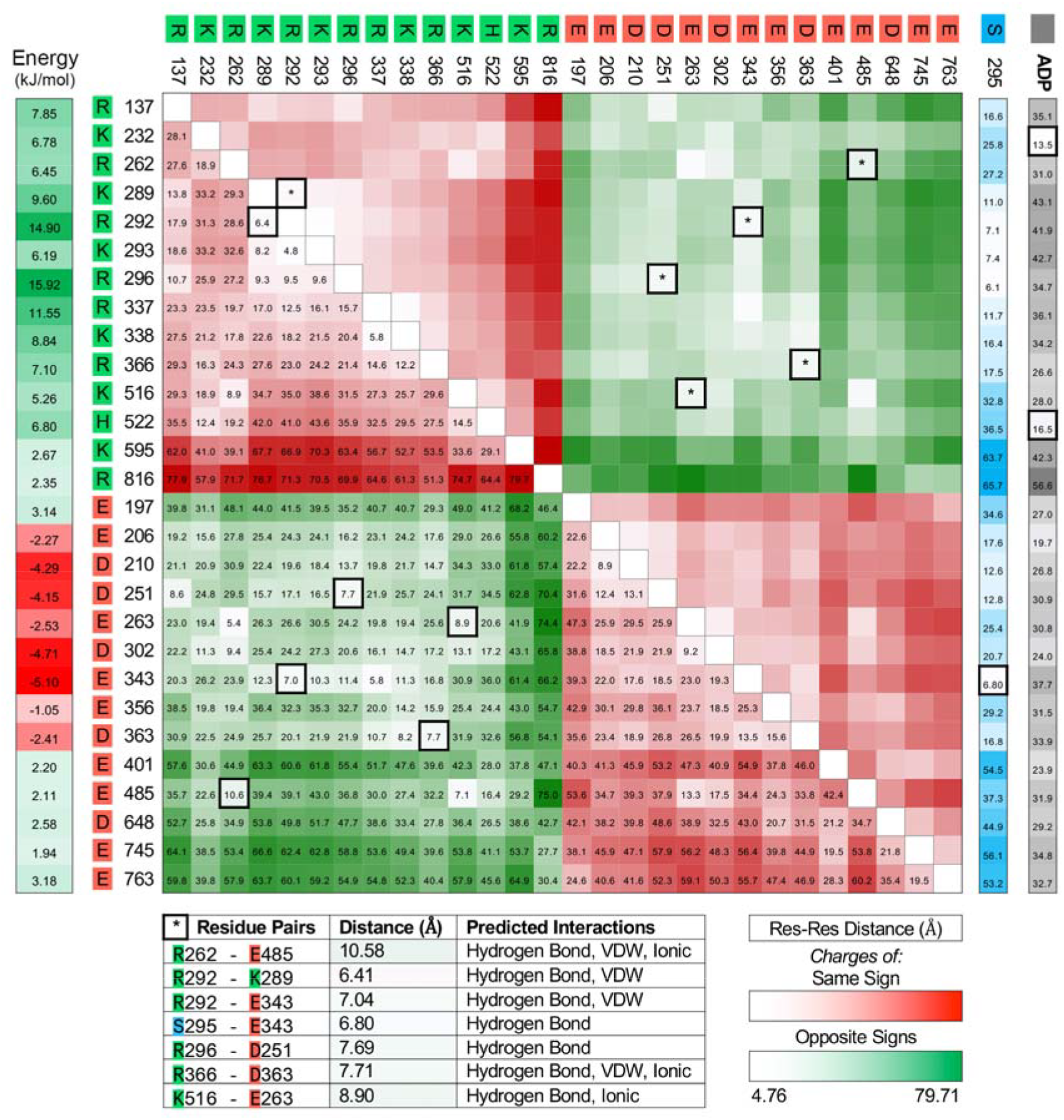
Inter-residue Distances for Significant Contributors to the Δ(Wq-q) between Unphosphorylated and Phosphorylated NLRP3 Models. Atomic distances and predicted interactions were calculated with DISTEVAL and RINGv2.0, respectively. RING interactions were defined as: hydrogen bonds, if the distance between acceptor and donor atoms was ≤ 3.5 Å and the hydrogen-donor-acceptor angle was ≤ 63°; van der Waals interactions, for residues with distances between atom surfaces of ≤ 0.5 Å; or salt-bridges, if the distance between the mass centers of oppositely charged groups was ≤ 4.0 Å. Residues were sorted by number, then charge (single letter amino acid code in bright red: negative, in bright green: positive). Energy values for every residue are listed to the left of the heat-map and are coloured from most positive (green) to most negative (red). Distances are coloured in a spectrum from white to green or white to red for residues of the same or opposite charges, respectively (Res-Res distances). Additionally, residue distances (Å) from S295 (blue) and ADP (gray) are displayed. If an interaction was reported by RING-2.0, the matrix value is outlined with a thick black border and an asterisk on the top half. Specific interactions between residue pairs are listed below the matrix.

## CONCLUSION

MD simulations identified specific attraction and repulsion events associated with Ser295 phosphorylation-induced conformational changes of the NACHT domain (**Figures 1 and 2**). It seems plausible that the local structural transformations could have consequences on the catalytic efficiency of the enzyme and suppress nucleotide turnover within the NBD. The change in NBD topology may impede the opening mechanism observed between the NACHT and LRR domains of NLRP3 in the unphosphorylated MD simulation ^17^. This is in accordance with established data demonstrating the inhibitory effect of Ser295 phosphorylation on NLRP3 prior to activation and even the disassembly of activated NLRP3 inflammasomes following PKA stimulation and phosphorylation of Ser295 ^13,30^. Alternatively, the inhibitory effect of pSer295 could result from conformational impacts on other regions distal to the core motifs of the ATP-binding pocket. This could include the homotypic interaction interface in the PYD domain for the adaptor molecule ASC ^31^, or the interface for NEK7 that involves the LRR and NACHT domains ^20,29^; both are critical for inflammasome assembly and activation ^2^. The results are unable to explain the contrasting impact of Ser295 phosphorylation by PKA (inhibitory) ^13,30^ and PKD (activating) ^16,32^. However, the global conformational changes in NLRP3 structure induced by Ser295 phosphorylation could also provide a mechanism to enable dissociation of NLRP3 from mitochondria-associated membranes and allow for cytoplasmic inflammasome assembly. Now that several x-ray crystal and cryo-EM structures of active and inactive NLRP3 are available, the collection of empirical structural data for phosphorylated forms of NLRP3 represents the next unmet challenge that will advance understanding of this important immune signaling platform.

## ACKNOWLEDGEMENTS

This research was supported by a Discovery Grant from the Natural Sciences and Engineering Research Council of Canada (NSERC; RGPIN/04379-2019 to JAM). C.F.S. was supported by an NSERC Postgraduate Doctoral Scholarship.

## AUTHOR CONTRIBUTIONS

All persons designated as authors qualify for authorship, and all those who qualify for authorship are listed. C.F.S. designed and conducted the experiments, completed data analyses, prepared figures, and co-wrote the manuscript. J.A.M. coordinated the study, provided trainee supervision, co-wrote the manuscript, and made intellectual contributions to the project. Both authors approved the final version of the manuscript and agree to be accountable for all aspects of the work in ensuring that questions related to the accuracy or integrity of any part of the work are appropriately investigated and resolved.

